# Decoding Glycomics: Differential Expression Reimagined

**DOI:** 10.1101/2023.08.04.551938

**Authors:** Jon Lundstrøm, James Urban, Daniel Bojar

**Affiliations:** Department of Chemistry and Molecular Biology, University of Gothenburg, Gothenburg, Sweden. Wallenberg Centre for Molecular and Translational Medicine, University of Gothenburg, Gothenburg, Sweden

## Abstract

Glycomics, the comprehensive study of all glycan structures in a sample, is a rapidly expanding field with substantial relevance for understanding physiology and disease mechanisms. However, the complexity of glycan structures and glycomics data interpretation present significant challenges, especially when it comes to differential expression analysis. Here, we present a novel computational framework for differential glycomics expression analysis. Our methodology encompasses specialized and domain-informed methods for data normalization and imputation, glycan motif extraction and quantification, differential expression analysis, motif enrichment analysis, time series analysis, and meta-analytic capabilities, allowing for synthesizing results across multiple studies. All methods are integrated into our open-source glycowork package, facilitating performant workflows and user-friendly access. We demonstrate these methods using dedicated simulations and various glycomics datasets. Our rigorous approach allows for more robust, reliable, and comprehensive differential expression analyses in glycomics, contributing to the advancement of glycomics research and its translation to clinical and diagnostic applications.

## Introduction

Glycomics, predominantly assessed via mass spectrometry^1^, plays a crucial role in understanding numerous biological processes in which glycans or complex carbohydrates are involved, such as disease pathogenesis^2, 3^. Glycans, for instance linked to proteins or lipids, are involved in critical biological events such as cell signaling, the immune response, and pathogen-host interactions^2^, and can interact with proteins via specific substructures, such as the Lewis antigens^4^. Besides functional importance, characteristic structural differences also exhibit substantial diagnostic potential, with elevated expression of sialyl-Lewis structures in various forms of cancer^5^, among others^6^. Therefore, precise elucidation of glycan structures and their differential expression across different conditions is paramount for both fundamental and clinical research.

However, differential expression analysis in glycomics presents significant computational and statistical challenges. The complexity of glycan structures, their variable expression levels, and the intricacy of high-dimensional glycomics data demand sophisticated analytical tools. Existing methods, while making substantial contributions^7, 8^, often fall short in addressing the complex nature of glycomics data in all its facets, particularly in dealing with missing data, normalizing data variance, and accurately determining differential expression with high sensitivity. Further, the current state-of-the-art of ad-hoc choosing relevant glycan substructures or features for differential expression testing, coupled with the frequent lack of multiple testing correction, renders results across studies incommensurable and has a high chance to result in p-hacking, even unconsciously.

Previous approaches to address this gap in the field have served to highlight the importance of properly analyzing glycomics data at various levels^8, 9^, yet have not found widespread adoption in the field and could be further improved by adopting state-of-the-art techniques from differential expression analysis in proteomics and metabolomics, as well as glycomics-specific advances.

Next to methodological drawbacks in how glycomics data are typically analyzed, the field also lacks various domain-adapted workflows for non-standard data types, which have been implemented in transcriptomics, proteomics, and/or metabolomics. These include time-series analyses, multivariate analyses, multi-group analyses, and meta-analyses. Specifically, we envision that glycomics has a need for analyzing various types of data on the sequence, motif, and motif set level, which needs to be flexibly incorporated, user-friendly, and fueled by state-of-the-art statistical robustness and sensitivity, given the heterogeneous nature of glycomics data and the difficulty of acquiring large datasets.

In light of these challenges, we developed a comprehensive computational framework that could provide enhanced capabilities for differential glycomics expression analysis and fill this gap. Our platform integrates various data normalization methods, sophisticated imputation methods for missing data, (multivariate) differential expression analysis, time-series data analysis, ANOVA, and meta-analysis workflows. All introduced methods can be flexibly used with sequences and motifs, benefiting from the fully automated motif annotation and quantification workflow that we substantially improved in this work. All methods and workflows are fully integrated into glycowork version 0.8, with user-friendly wrapper functions. This integration makes highly optimized workflows possible, which take seconds to run on a normal laptop.

Throughout this work, we use carefully simulated glycomics data, to evaluate the steps of our workflows, as well as real-world glycomics data to demonstrate that insights can be gained rapidly and even from relatively modest sample sizes. In the course of doing so, we uncover numerous new findings from reanalyzing glycomics data, which are also more reliable thanks to our normalizations and multiple testing corrections. The objective of this study is to provide more reliable, comparable, and comprehensive tools to facilitate advanced glycomics research, aiding in the translation of glycomic insights into diagnostic and therapeutic strategies.

## Results

### Automating glycan motif quantification at scale

Given that glycan motifs often drive glycome properties and functions, for instance by binding proteins in a modular manner^10^, analyzing glycomics data on a substructure level might offer benefits in interpretation. This is especially true as the same motif might be present in several glycans, presenting a confounding element on a sequence level. Thus, motif analysis combines information from several, biosynthetically related, glycans. Related work has shown that this indeed leads to a substantial increase in statistical power, compared to analyzing full sequences^9^.

Our glycowork Python package already contained several workflows for automatically annotating glycan motifs^11^. However, for the work presented here and included in glycowork version 0.8, we have completely refactored our approach to motif quantification to be more performant, comprehensive, and proportional (Fig. 1).

**Figure 1.**
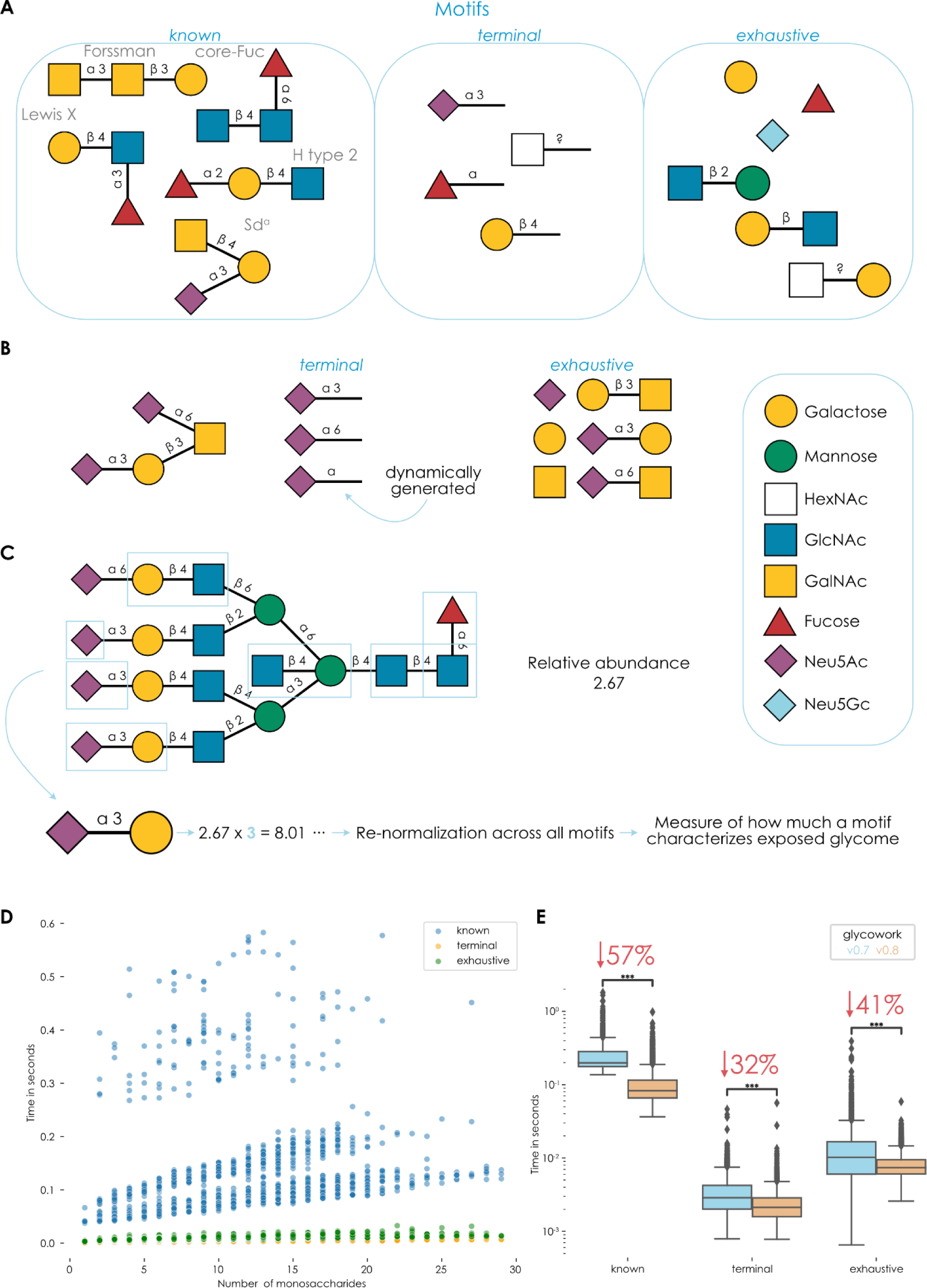
A reworked motif annotation and quantification platform within glycowork. **A)** Overview of the types of standardized motifs that are generated within glycowork. The different motif groups are categorized by their keyword that can be used in many functions within glycowork, from *annotate_glycan* to the *get_differential_expression* workflow described below. **B)** Illustration of the dynamic generation of useful motifs beyond provided sequences. Annotation functions within glycowork will automatically generate more general motifs (e.g., Neu5Acα2-?) and retain them if they capture non-identical information from their fully specified counterparts, allowing for easy enrichment tests. **C)** Proportional motif quantification. Glycan motifs (exemplified by blue boxes) are annotated, counted, and used as scaling factors to transform initial relative abundances into indicators of what proportion of the cellular surface is represented by this motif. **D)** Timing the motif annotation functions on physiological glycans. For all 4,260 human glycans up to 30 monosaccharides in size that are stored within glycowork (version 0.8), we timed the respective *annotate_dataset* portion for each feature_set option (“known”, “terminal”, “exhaustive”) and displayed the results for all those glycans as scatterplots. Differences in the distribution of “known” glycans are dependent on the motif density of a glycan, as more motifs imply more (expensive) subgraph isomorphisms tests. Timing tests were performed on an Intel(R) Xeon(R) CPU @ 2.20GHz. **E)** Comparing motif annotation speed between glycowork version 0.7 and 0.8. Similar to (D), the annotation functions were timed for all human glycans and compared between the current glycowork version and the one introduced in this work. Results are shown here via boxplots. Data are depicted as mean values, with box edges indicating quartiles and whiskers indicating the remaining data distribution. Note the logarithmic y-axis. Mean differences were tested with two-tailed Welch’s t-tests followed by Benjamini-Hochberg correction. The average percent decrease in runtime is shown for each keyword. All glycan structures or motifs in this work are depicted via the Symbol Nomenclature for Glycans (SNFG) and were drawn using GlycoDraw^12^. ***, p < 0.001; **, p < 0.01; *, p < 0.05; n.s., p > 0.05.

For this, we aimed for a standardized vocabulary of assayed glycan motifs, which still retained flexibility for new sequences as well as various degrees of structural annotation (e.g., uncertain linkages etc.). Therefore, our motif annotation pipeline contained three keywords, which can be mixed-and-matched at will (Fig. 1A). A set of 154 manually curated and named motifs (e.g., “LewisX” or “Sda”) can be accessed under the “known” keyword, while the keywords “terminal” and “exhaustive” generate motifs based on the provided sequences. The former will catalogue all occurring non-reducing end monosaccharides and their linkages, while the latter will count all mono- and disaccharides within a glycan. It should be noted that any motif overlap that may occur when keywords are combined is automatically removed within the annotation functions (see STAR Methods). These keywords placed an emphasis on terminal structures and smaller, modular substructures, which are commonly viewed as functional determinants within glycans^13^.

We realized that, while potent for mediating standardized analysis of differentially expressed substructures that indicated functional consequences, this workflow was occasionally subject to the limitations of nomenclature. If sialylation in general increased, this effect would be split between Neu5Acα2-3 and Neu5Acα2-6 in the case of analyzing glycans with the “terminal” keyword. Therefore, we implemented a system (Fig. 1B) that dynamically would generate the subsuming category (e.g., Neu5Acα2-?) and only retained it if it was not collinear with either specified version (i.e., if it contributed additional information), even if Neu5Acα2-? itself was not present in any glycan. This allowed us to probe more general trends in datasets.

If glycan substructures or features are further analyzed in glycomics data, the usual approach is to binarize information. This results in, for instance, analyzing “fucosylated” glycans (i.e., glycans with >0 Fuc vs glycans with 0 Fuc). This, however, loses valuable information about motif density. At equal concentration, an *N*-glycan exposing multiple antennae capped with Neu5Acα2-3 shapes the properties of the cellular surface more than a glycan with only one antenna that contains Neu5Acα2-3. We therefore developed a weighting scheme (Fig. 1C) that would count each motif (from the keyword classes mentioned above), retrieve the relative abundances of the associated glycans, and scale these abundances by the motif count. Followed by an additional normalization across all motifs, this then translated into an indication of the dominance or proportion of a given glycan motif across the whole glycome and is a better indicator for biologically relevant changes upon differential expression.

As for any analytical workflow, scalability is important. This will become especially relevant once glycomics is increasingly employed for sample characterization^14^, for instance aided by methods to automate data analysis^15^. Further, we aimed for timescales that would allow for (i) dynamic motif calculations, without any need for any pre-calculated intermediate results, and (ii) rapid re-running of workflows with different parameters (e.g., different motif keywords), facilitating interactive work. We therefore extensively optimized our motif annotation platform to fully annotate even larger glycans in milliseconds on a regular CPU (Fig. 1D). On average, this makes motif annotation in glycowork version 0.8 nearly twice as fast as in version 0.7 (Fig. 1E), despite it being more comprehensive and accurate in the new version. It should be noted that this also translated into gains that transcended the differential expression platform we present here and, for instance, eased the visualization of motif distributions via heatmaps^16^, the generation of biosynthetic networks^17^, and various other motif-level workflows that are enabled by glycowork.

### Establishing a robust and sensitive workflow for differential glycomics expression

Regardless of whether researchers are interested in the expression of glycan motifs or full sequences, several common issues arise. These include data sparsity, missing data, and heterogenous data, which inflates the false-negative rate of current differential glycomics expression analyses. This is paired with the current, all-too-common, practice of using unadjusted p-values to evaluate results, which in turn inflates false-positive rates. We thus set out to conceptualize a workflow that addressed these common challenges, still accommodated motif and sequence analysis, and balanced rigorous statistical procedures with the sensitivity needed for analyzing glycomics data.

On a conceptual level, this meant that we needed to implement robust methods for (i) data imputation, (ii) data normalization, (iii) statistical testing, and (iv) multiple testing correction, all of which had to be compatible with analysis on the sequence, motif, and motif set level (i.e., univariate and multivariate statistics). We note that, next to these features, we also implemented effect sizes (Cohen’s *d* / Cohen’s *d_z_* for univariate and Mahalanobis distance for multivariate statistics) as well as Levene’s test to test the homogeneity of variance across groups. In the following, we will discuss the choice of methods for each of these elements, which resulted in a performant workflow for engaging in differential glycomics expression analysis (Fig. 2A, Fig. S2) that we showcase with various examples throughout this work.

**Figure 2.**
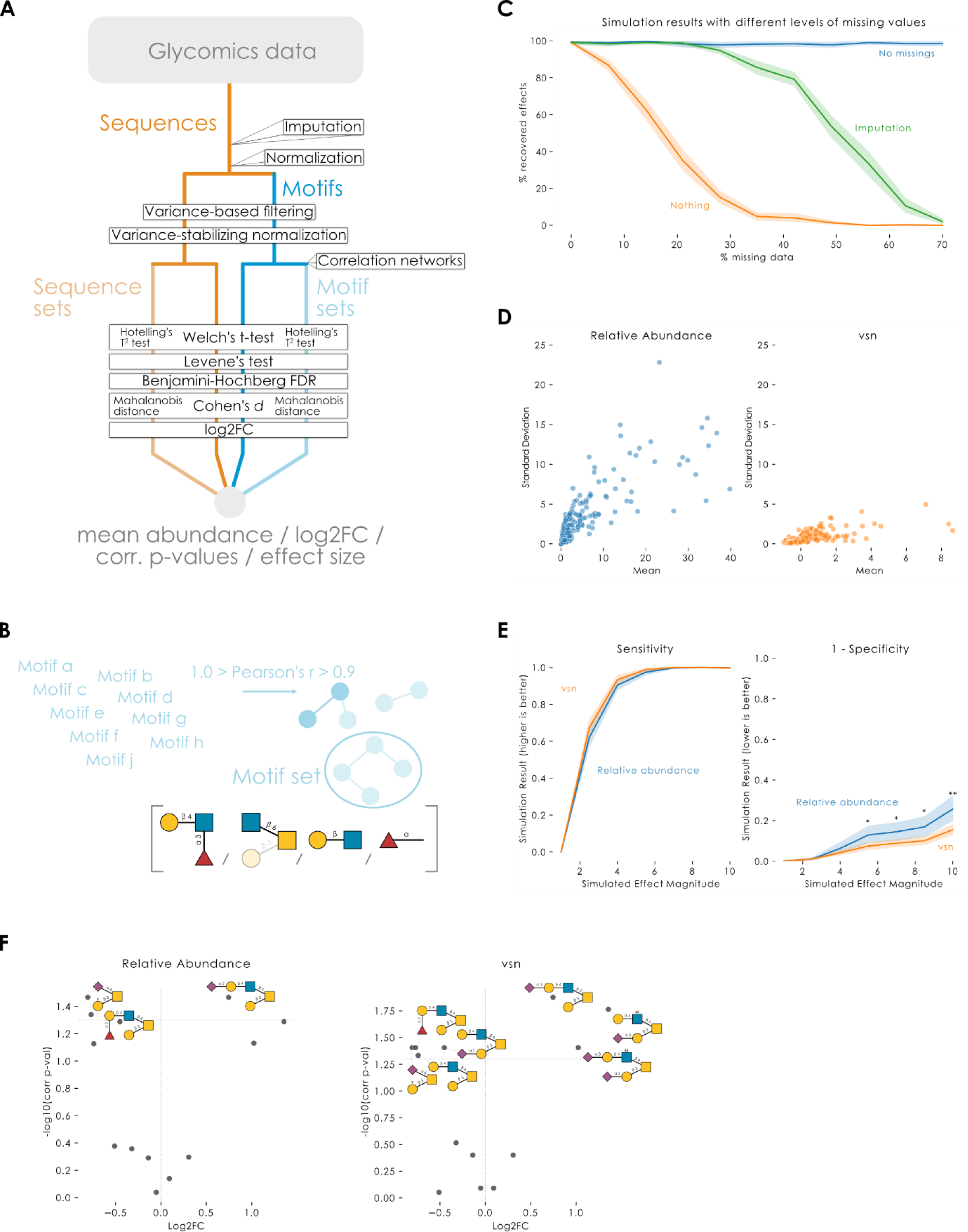
Establishing a statistically robust and sensitive workflow for differential glycomics expression analysis. **A)** Schematic overview of the *get_differential_expression* in glycowork (version 0.8). **B)** Overview of the construction of motif sets. If sets = True, *get_differential_expression* will aim to form sets of highly correlated sequences or motifs. Briefly, sequences or motifs with a high, but not perfect, correlation are connected in a graph and connected components of this graph constitute sets. These in-situ obtained correlated sets are then handled as multivariate tests within *get_differential_expression* and constitute a form of enrichment analysis. **C)** Glycomics data imputation makes the workflow robust to missing data. From 0 to 70% of randomly missing data, we sampled 128 glycans from a Dirichlet distribution (see STAR Methods for details) for 50 experiments per level of missing data, for two groups of ten replicates each (N = 1000 for each level of missing data). Whether the simulated effects were correctly identified was tested for the workflow with and without data imputation, including a comparison group without missing data. **D)** Variance stabilizing normalization (vsn) minimizes heteroscedasticity. Glycan abundances from nine experimental datasets^18–26^ were first normalized by dividing each abundance by the total abundance of the sample (i.e., relative abundance). Then, vsn was performed by log-transformation of the data and standard scaling to zero mean and unit variance, using the *variance_stabilization* function in glycowork. **E)** Variance stabilizing normalization improves workflow specificity. Simulations proceeded similar to (C), with the main difference being that no data were missing. Instead, the simulated effect magnitude was varied by the concentration parameter when sampling from the Dirichlet distribution, scaling from 1 to 10 in seven evenly spaced steps. Statistical significance was established with two-tailed Welch’s t-tests and the resulting p-values were adjusted using the Benjamini-Hochberg method. **, p < 0.01; *, p < 0.05. **F)** *O*-glycomics data from FFPE tissue of healthy and basal cell carcinoma (BCC) samples^26^ were analyzed with the *get_differential_expression* function to illustrate the impact of vsn-transformation of relative abundance values. Results are depicted as volcano plots obtained via the *get_volcano* and *annotate_figure* functions.

In addition to sequence and motif level analysis, we also wanted to support the analysis of motif sets. An idea that originated in metabolomics analysis^27^, set analysis assembles highly correlated metabolites to compare them as a group, via multivariate statistics, across conditions, reasoning that a common biological process stands behind their differential expression. As glycan motifs can have strong biosynthetic relationships (e.g., Sialyl-Lewis X contains a Lewis X motif as well as a Neu5Acα2-3 motif), we hypothesized this to be a valuable addition to enhance sensitivity. Practically, inspired by metabolomics^27^, we automatically constructed sequence or motif sets via correlation networks, where two entities with a (positive) correlation between 0.9 and 1 (excluding 1) were connected via an edge (Fig. 2B). Each connected component within such a network constituted a set that was compared via Hotelling’s T^2^ test (a multivariate extension of the t-test^28^), with the Mahalanobis distance as an estimate of the multivariate effect size^29^.

A recent analysis of imputation strategies for proteomics data has identified MissForest^30^, a Random Forest-based approach, as the current best-in-class method for data imputation^31^. We thus implemented a version of MissForest within glycowork to perform our data imputation. This machine learning-based imputation strategy, in contrast to single-value imputations (e.g., replacing all missing values with 0.1) that are commonly used in glycomics data, does not profoundly affect the underlying distribution of glycan abundances and thus should be more robust to artifacts. To test the impact of missing data on the workflow, with and without imputation, we generated simulated glycomics data with 0 to 70% of randomly missing values (see STAR Methods for details). At 30% missing values, differential expression with our data imputation approach recovered the simulated effects with a sensitivity comparable to analysis of the dataset without missing values. In contrast, employing no imputation decreased the sensitivity to below 20% (Fig. 2B).

Glycomics abundance data are inherently heteroscedastic, displaying increasing variance across the range of measured abundances (Fig. 2D). In order to minimize this phenomenon, and increase statistical power, we performed variance stabilizing normalization (vsn) by log-transforming the data followed by standard scaling to zero mean and unit variance (Fig. 2D). Using simulated glycomics data, we compared the workflow performance between using relative abundances and the corresponding vsn-transformed values. Here, we observed a significantly lower false-positive rate (1 – specificity) at higher simulated effect magnitudes upon vsn transformation (Fig. 2E). We further tested the impact of vsn transformation on the workflow performance by reanalyzing *O*-glycomics data from formalin-fixed paraffin-embedded (FFPE) tissue collected from 20 patients with basal cell carcinoma (BCC)^26^.

Differential expression analysis using relative abundances as the input yielded three significantly changed structures, reflecting a general shift from core 1 to core 2 *O*-glycans as originally reported^26^. Vsn transformation further enabled the detection of four additional significantly changed glycans, showing an increase in sialylated, sulfated core 2 *O*-glycans, as well as downregulation of their respective biosynthetic precursors (Fig. 2E).

At the core of the differential expression testing, we chose to use two-tailed Welch’s t-tests (Fig. 2A) for their robust empirical performance with small sample sizes and their consideration of potential heteroscedascity in the input data^32^. Contrary to common misconceptions, the various t-tests do not assume normally distributed input data but rather normally distributed differences in means (for example), which is obtained even with very small samples^33, 34^. We still made sure that this assumption holds true in glycomics data as well and compared it with a nonparametric test (Fig. S1), which showed the equivalence of our approach and highlighted the slightly lower rate of false-positives that can often be achieved with parametric tests^34^. As further discussed below, an even better approach to this might lie in generalized linear models (GLMs) and related methods^35^, yet given typical sample sizes and variances observed today, we believe that our approach is a more fruitful compromise until future technologies will unlock this realm for glycomics data.

In order to showcase the sensitivity of motif-level analysis, we modified the data simulation approach to scale the abundance of all glycans containing a particular motif of interest. Across randomly sampled human *O*-glycans, we scaled the concentration parameter of glycans containing ‘Neu5Acα2-6’ by 1.25 in the test condition, reflecting a modest log2FC of ∼0.3. As expected, no significantly differentially expressed glycans were detected by a sequence-level analysis, while in contrast, the motif-level analysis correctly identified the terminal motif ‘Neu5Acα2-6’ as significantly regulated (Fig. 3A). This also made abundantly clear that motif-level effects can be entirely missed, when analyzing glycomics data only on the sequence-level.

**Figure 3.**
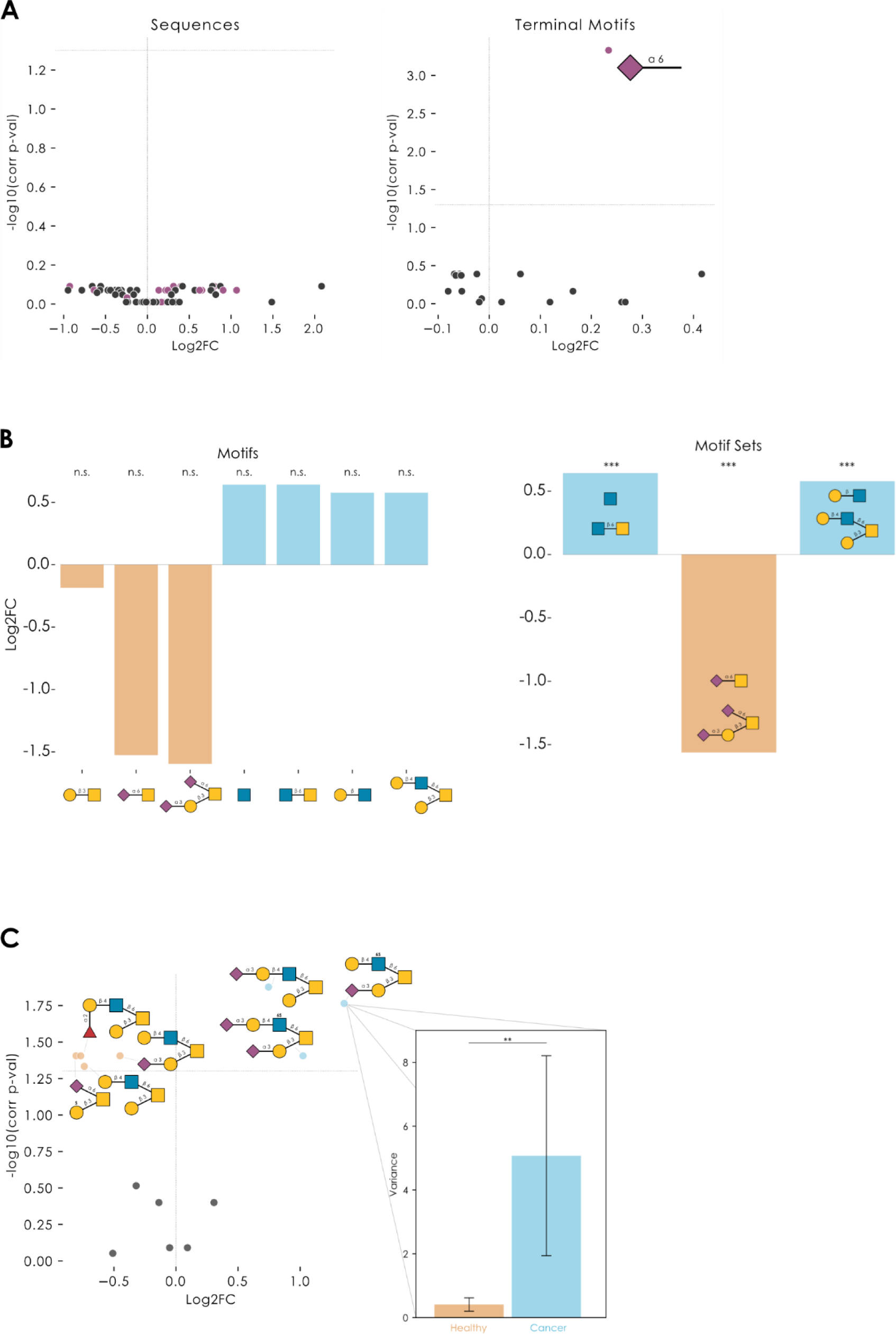
Analyzing differential glycomics expression with a performant workflow. **A)** Volcano plots comparing the workflow sensitivity at the sequence and motif level. In the reference condition, random human *O*-glycans from the dataset within glycowork were assigned to simulated abundances. In the test condition, concentration parameters of structures containing ‘Neu5Acα2-6’ were multiplied by 1.25 (log2FC ∼0.3). Sequences or motifs containing ‘Neu5Acα2-6’ are labeled in purple. N = 5 per condition. **B)** The *O*-glycomics data from prostate cancer samples^25^ were analyzed with *get_differential_expression* at the motif level, either with sets = False or sets = True, to illustrate the enhanced sensitivity achieved by analyzing sets of highly correlated motifs. For sets = False, the seven motifs with the lowest adjusted p-values are shown, together with their log2-transformed fold changes (cancer versus healthy). **C)** The *O*-glycomics data from the basal cell carcinoma dataset from Möginger et al.^26^ were analyzed with *get_differential_expression* at the sequence level, followed by a visualization of the significantly differentially expressed glycans via the *get_volcano* and *annotate_figure* functions of glycowork. Shown as an inset are the variances of healthy and cancer samples for the only glycan for which Levene’s test for Equality of Variances yielded a significant p_adj_. The bar graph shows the variances, together with a 95% confidence interval estimated by bootstrapping 1,000 times, using sampling with replacement. ***, p < 0.001; **, p < 0.01; *, p < 0.05; n.s., p > 0.05.

To ensure that our motif annotation and quantification approach was performant, we compared motif abundances derived from GlyCompare^9^ with those we obtained within glycowork on the abovementioned BCC dataset^26^ (Fig. S3, Supplementary Table 3). While GlyCompare-features did recapitulate the increase in Neu5Acα2-3, glycowork-derived motif abundances yielded a greater number of differentially expressed motifs that reached statistical significance within our workflow. Here, we also want to emphasize our end-to-end approach of dynamically quantifying motifs within the analysis, thereby making any pre-processing or post-processing steps unnecessary, which is not the case for most alternative approaches. A loaded table of relative abundances of glycans can thus, quite literally, be analyzed with a single line of code. This makes our approach inherently more scalable and usable.

As mentioned before, our approach of optionally analyzing highly correlated sets of glycan motifs via correlation networks, as pioneered in metabolomics^27^, substantially enhanced sensitivity and robustness to differences in data annotation (such as linkage uncertainties in some structures). As an example, we applied this to a dataset of 55 prostate cancer *O*-glycomics samples^25^, in which a motif-level differential expression analysis did not yield any significant differences between tumor and healthy samples after multiple testing correction (lowest p_adj_: 0.07; Fig. 3B). In contrast, motif set-level analysis revealed biosynthetic clusters of glycan motifs that were significantly dysregulated in cancer, such as a lower level of extended Neu5Acα2-6GalNAc containing glycans and higher levels of extended core 2 *O*-glycans (Fig. 3B). This showcased the benefits of motif set-level analysis for, often heterogeneous, glycomics data.

Most statistical tests investigate whether samples differ in their mean values for entities such as glycans. However, it is a well-known fact that diseases, such as cancer, also manifest with a substantial degree of heterogeneity^36^. Especially bulk approaches, such as glycomics, can be affected by this heterogeneity. Our workflow therefore also contained Levene’s test for Equality of Variances, which, when significant, indicates that variances are significantly different across the comparison groups. It is important to note that this can occur even in the absence of significant mean differences. Applied to an example dataset of *O*-glycomics data of basal cell carcinoma samples and healthy controls^26^ (Fig. 3C), we not only found significant mean differences (sulfated/sialylated glycans upregulated; neutral/fucosylated glycans downregulated) but also one sequence, Neu5Acα2-3Galβ1-3(Galβ1-4GlcNAc6Sβ1-6)GalNAc, that exhibited a highly significant Levene’s test. This supported our choice of Welch’s t-test, as the regular t-test would not be warranted in such a case of heteroscedasticity. This increased variance of specific sequences in, for instance, heterogeneous diseases such as cancer^36^ could then indicate either increased noise in the regulation of responsible enzymes or cell type subgroups within the bulk sample which might be identifiable via these glycans. In either case they are promising starting points for follow-up investigations.

### Accommodating various types of glycomics data and experimental designs

While comparing two groups regarding their glycome is an essential and almost ubiquitous task in analyzing glycomics data, we also wanted to support different data structures. A common problem occurs when more than two experimental groups are compared and the relevant pairwise differences are not known a priori. Here, the standard approach would be an analysis of variance (ANOVA), followed by post-hoc tests of glycans/motifs which show any significant differences between groups. We implemented this functionality in the *get_glycanova* function, which triggers Tukey’s HSD (honestly significant difference) test if any ANOVA result was significant. As this test controls the family-wise error rate, the final p-values are adjusted for multiple testing. Apart from the statistical testing itself, the *get_glycanova* function benefited from all improvements made to the *get_differential_expression* function, such as our data imputation strategy or variance-stabilizing normalization.

To illustrate this, we used a previously reported glycosphingolipid dataset^37^, which investigated ganglioside expression from different tissues across various kinds of ceramide portions. Using the *get_glycanova* function, we probed whether any glycan motif was differentially expressed across the different pools of ceramides. This identified elevated motifs, such as the increased expression of GalNAcβ1-4Gal substructures (from the Sd^a^ motif) in short and long, but not middle-sized, ceramide portions (Fig. 4A).

**Figure 4.**
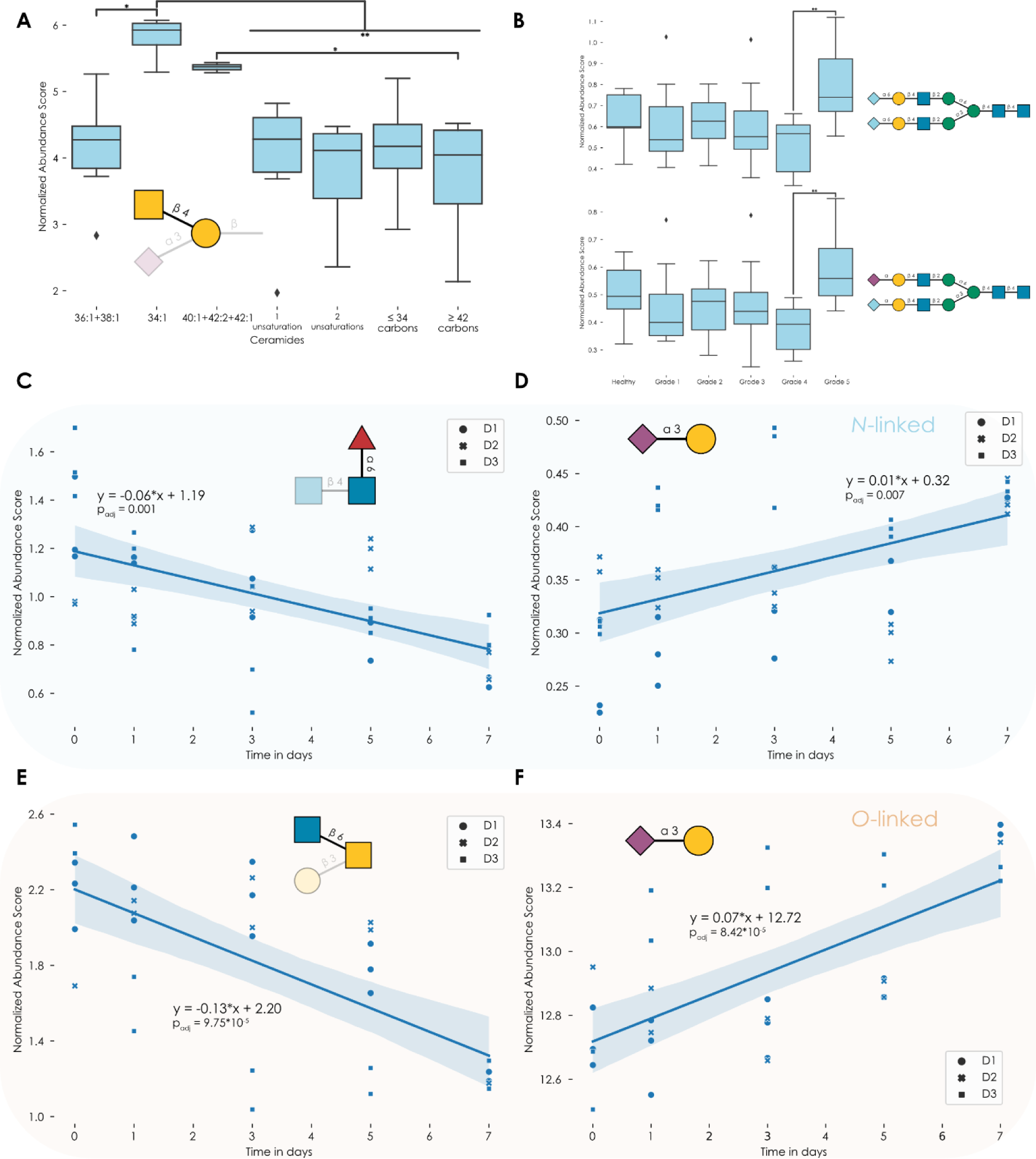
Analyzing multi-group data and glycomics time series data. **A-B)** We used the *get_glycanova* function for a motif analysis of the glycosphingolipid dataset in Sibille et al.^37^ (A) and a sequence analysis of the *N*-glycan dataset in Kawahara et al.^25^ (B). Asking the question whether any glycan motif/sequence exhibited differential expression across the different types of ceramide (A) or prostate cancer grades (B), the *get_glycanova* function engages in an ANOVA and pairwise Tukey’s HSD tests for any significant results. The example of GalNAcβ1-4Gal (A), as part of the Sd^a^ motif, and two Neu5Gc-containing *N*-glycans (B) are shown here via boxplots. Data are depicted as mean values, with box edges indicating quartiles and whiskers indicating the remaining data distribution. **C-F)** Representative motifs showing significant changes in *N*-linked (C-D) or *O*-linked glycans (E-F). For the data reported by Hinneburg et al.^38^, we used the *get_time_series* function from glycowork, with motifs = True, for both *N*- and *O*-glycomes. From the full table of results (Supplementary Table 4), we chose the core fucose motif (C), terminal α2-3 linked sialylation in *N*-linked glycans (D), the core 2 *O*-glycan determinant (E), and terminal α2-3 linked sialylation in *O*-linked glycans (F) as representative examples. Shown are scatter plots of the respective normalized abundance scores calculated by the *quantify_motifs* function, with different shapes for points originating from different donors. Overlayed is a fitted linear regression line, including the 95% confidence interval band, and the regression equation as well as the Benjamini-Hochberg-adjusted p-value of the regression coefficient, derived from a t-test. ***, p < 0.001; **, p < 0.01; *, p < 0.05; n.s., p > 0.05.

Of course this workflow can similarly be applied on the glycan sequence level, and we used this in a glycomics-adapted ANOVA analysis of an *N*-glycomics dataset of prostate cancer patients^25^ to, for instance, show that several complex *N*-glycans were differentially expressed only between grades 4 and 5 in prostate cancer (Fig. 4B), which could indicate their usefulness in patient stratification.

Another example of glycomics data structures would be time-series data, a common setup to test the effect of a drug or monitor a cell differentiation program. While the analysis of such data can be, and often is, reframed as a group comparison (e.g., differential expression of first timepoint vs last timepoint), such an approach lowers sensitivity and may miss transient expression changes.

Instead, in the *get_time_series* function, we chose an approach that fitted an Ordinary Least Squares (OLS) regression to each glycan or motif and performed t-tests to evaluate whether the regression coefficients were significantly different from zero. We are aware that this approach may underestimate non-linear trajectories and other effects but consider it a compromise of currently available data quantities, and an improvement compared to current practices. For enhanced functionality and comparability with the methods presented above, the *get_time_series* function used the same data imputation, normalization, motif quantification, and multiple testing correction strategies employed in our other functions.

To illustrate the superior sensitivity and usefulness of this analytic approach, we chose an available glycomics dataset from the literature, in which the differentiation of human monocytes to macrophages was monitored by *N*- and *O*-glycomics time-series data^38^. Analyzing these two time-series datasets with our workflow resulted in a large number of significantly changing glycan sequences and motifs during this process (Fig. 4C-F, Supplementary Table 4), despite our more rigorous workflow that included multiple testing correction.

One of the most prominent results for *N*-linked glycans was the decrease of core fucosylated structures over time (Fig. 4C), also noted by the authors of the original study. However, the much lower p-value, compared to the t-test comparing day 0 and day 7 from Hinneburg et al., clearly showcased the increased sensitivity of our approach. Therefore, while the original authors did report no significant changes in *N*-glycan sialylation, we also noted a significant increase in, specifically α2-3-linked, Neu5Ac among *N*-glycans (Fig. 4D).

For *O*-linked glycans, we first recapitulated the major finding of Hinneburg et al., a decrease in core 2-containing structures (Fig. 4E). We then went on and further demonstrated a significant increase in α2-3-linked Neu5Ac (Fig. 4F), mirroring its increase in *N*-glycans and hinting at a general upregulation of this motif in human macrophages upon differentiation.

We recognize that non-linear expression trajectories might well be possible and relevant for glycomics data and thus also allow for investigating polynomial fits to time series data with the “degree” keyword argument to *get_time_series*. We demonstrated the usefulness of this with data from free milk oligosaccharides^39^ (Fig. S4), which often exhibit non-linear trajectories throughout lactation that can be better captured with polynomial functions. Future work could increase the repertoire of *get_time_series* further, to accommodate more complex temporal expression patterns.

### Applying the differential expression workflow to a cancer *O*-glycomics meta-analysis

On the one hand, substantial heterogeneity in glycan expression and different measurement techniques can make cross-study comparisons hard. Yet, on the other hand, increasing data collection capabilities have resulted in a growing set of glycomics datasets targeting the same underlying disease, such as different forms of cancer. Following more mature fields, we hypothesized that combining results from different studies and different patient cohorts could thus allow for more reliable estimates of conserved changes in a disease glycome. The state-of-the-art approach for this procedure is encompassed in the method family of meta-analyses, in which systematic literature searches are followed by procedures to combine effect sizes across studies.

To accommodate this growing potential in the field of glycomics, we have implemented fixed-effects and random-effects meta-analysis^40^ capabilities into glycowork version 0.8, which were designed to seamlessly work with the differential expression methods mentioned above and can be used to readily combine a series of dataset analyses with the *get_meta_analysis* function.

We then set out to demonstrate the potential of this functionality by performing an explorative mini meta-analysis of pan-cancer *O*-glycomics data from human patient samples (see STAR Methods for details), as there are already at least systematic review-type of analyses for pan-cancer *N*-glycomics data^41^, yet not for *O*-glycomics data. Surveying the current literature on such *O*-glycomics studies, in which the underlying data have been published as well (essential for re-analysis with our approach), resulted in nine publications with eleven cohorts (total N = 223) from a variety of cancer types.

While there are many invaluable reviews about *O*-glycan dysregulations in cancer^42–44^, these mostly comprise manual and subjective collations of various findings across the years. This is certainly valuable, yet lacks the technical rigor of an actual meta-analysis, especially as all the individual findings that led to these collective evaluations came from very heterogeneous data analysis approaches and, due to the lack of multiple testing correction, would not have been findings in many cases in rigorous analyses.

Analyzing effect sizes of differentially expressed glycans and glycan motifs across all these datasets allowed us to reveal which glycan sequences and motifs in fact were consistently dysregulated across the types of cancer included in this analysis (Supplementary Table 5). The most consistently dysregulated sequence in cancer was Neu5Acα2-3Galβ1-4GlcNAcβ1-6(Neu5Acα2-3Galβ1-3)GalNAc, always being upregulated in tumor samples, albeit with considerable heterogeneity in its expression (Fig. 5A). Overexpressed sequences such as this could, in the future, become general purpose cancer biomarkers as well as targets for recruiting anti-tumor proteins^45^. Importantly, structures such as this one (α2-3-sialylated core 2 *O*-glycans) have been recently identified to define cancer stem cell populations within breast cancer^46^.

**Figure 5.**
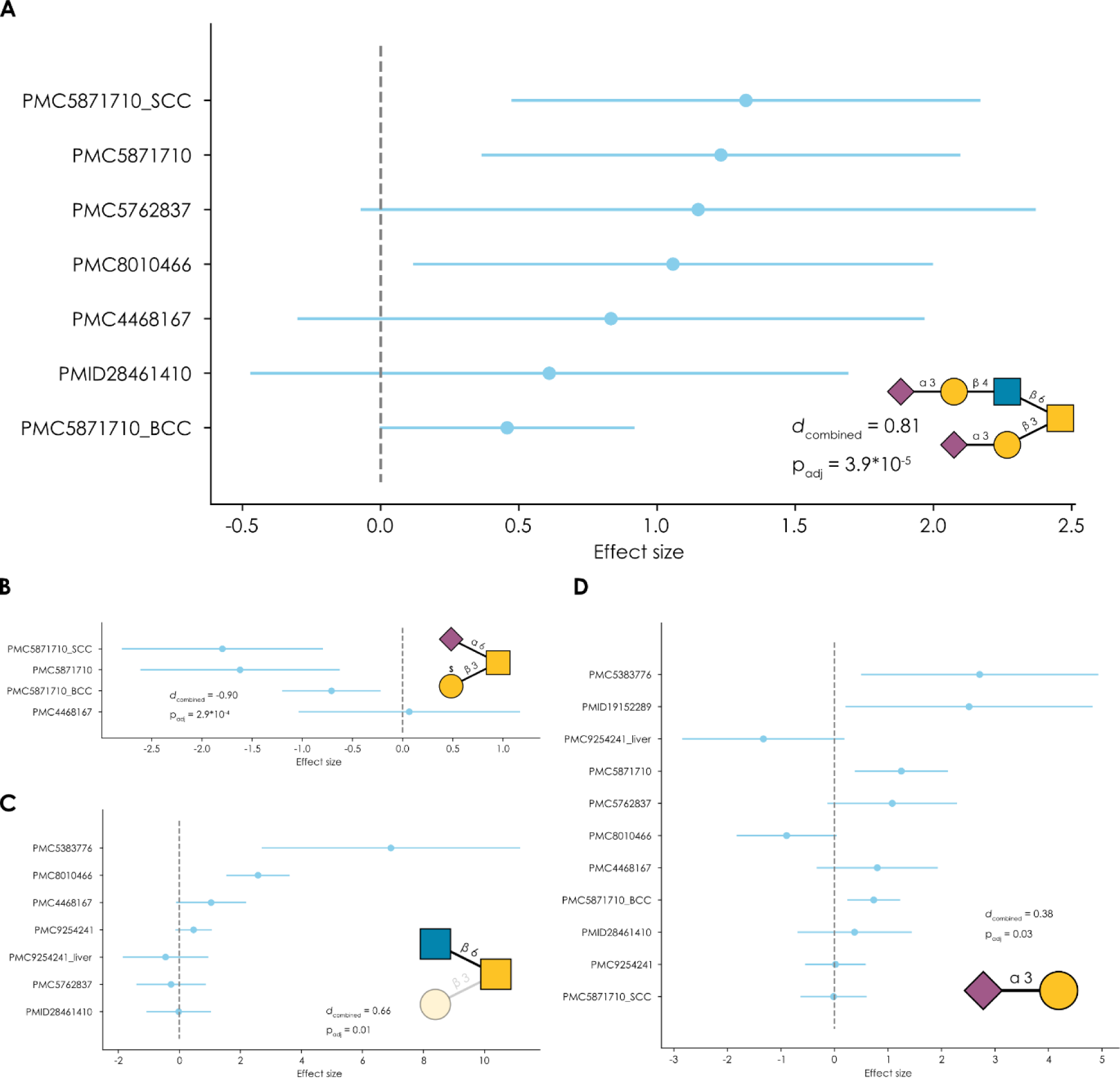
Meta-analysis of cancer *O*-glycomics data using the *get_meta_analysis* function in glycowork. **A-D)** For a total of 11 patient cohorts (from nine studies, total N = 223) of *O*-glycomics data from tumor and healthy tissue of gastric, skin, liver, prostate, colorectal, and ovarian cancer, we used the *get_differential_expression* function on the sequence (A-B) and motif (C-D) level to obtain effect sizes and effect size variances. Using a fixed-effects meta-analysis via the *get_meta_analysis* function, we obtained combined effect sizes (Cohen’s *d*) and adjusted p-values for each sequence or motif (Supplementary Table 5). Shown are Forest plots of representative sequences (Neu5Acα2-3Galβ1-4GlcNAcβ1-6(Neu5Acα2-3Galβ1-3)GalNAc (A), GalOSβ1-3(Neu5Acα2-6)GalNAc, (B)) or motifs (core 2 *O*-glycans (C), Neu5Acα2-3Gal (D)), also derived from the *get_meta_analysis* function. For each study recording this sequence or motif, the derived effect size and its 95% confidence interval is shown. In addition, the combined effect size and the associated adjusted p-values are also shown.

Conversely, other sequences such as GalOSβ1-3(Neu5Acα2-6)GalNAc were downregulated in all studies in which they were identified (Fig. 5B). It also speaks to a more general trend we have noticed throughout this work, that differentially expressed glycans often were relatively lowly abundant structures, which further hampers this detection due to greater relative variance. Importantly, however, this does not mean that we find sulfated glycans in general to be downregulated in the case of cancer, as other structures, such as Neu5Acα2-3Galβ1-3(Galβ1-4GlcNAc6Sβ1-6)GalNAc, were found to be significantly upregulated (*d*_combined_ = 0.6, p_adj_ = 0.04) and other researchers also identified specific upregulated sulfated *O*-glycans in cancer^47^.

Here again, an analysis of the underlying motifs might shed light on patterns of dysregulation that would not be apparent at the individual sequence level, such as the overexpression of core 2 *O*-glycans in cancer (Fig. 5C), which is a known finding^18, 26^. Another example of this can be found in sialylation. While Neu5Ac in general^48^, and the sialyl-Tn (Neu5Acα2-6GalNAc) sequence in particular^49^, are found to be upregulated in cancer, we find this picture to be somewhat more complicated.

While Neu5Acα2-3 is indeed widely upregulated in cancer *O*-glycomes (Fig. 5D), total Neu5Acα2-6 is actually downregulated, on average, in reported cancer glycomics data (Supplementary Table 5). The reason for this is twofold: (i) sialyl-Tn is often not reported in cancer *O*-glycomics data (only six of our 11 cohorts even report its expression in any sample, including several cohorts with basically identical mean values of sialyl-Tn across groups), and (ii) di-sialyl-T antigen (Neu5Acα2-3Galβ1-3(Neu5Acα2-6)GalNAc) was consistently downregulated in the reported datasets (*d*_combined_ = −0.5, p_adj_ = 0.01; Supplementary Table 5). While sialyl-Tn did exhibit a moderate upregulation (*d*_combined_ = 0.3), this did not reach statistical significance (p_adj_ = 0.68; Supplementary Table 5). As a consequence, the total pool of Neu5Acα2-6 containing glycans registered as downregulated on average. While we acknowledge that these structures are predominantly biosynthesized by different enzymes^50^ (ST6GALNAC1 for sialyl-Tn and ST6GALNAC3/4 for di-sialyl-T), the fact remains that the number of Neu5Acα2-6 epitopes that are exposed on the cell surface seems to decrease in cancer on average, shifting towards Neu5Acα2-3, at least according to currently published glycomics data. Insights such as these are important reasons for conducting such meta-analyses across the entire measured glycome and at the motif level.

Overall, we conclude that the current prevalence of available *O*-glycomics data from cancer samples is lower than commonly believed. Especially given (i) the high heterogeneity within and between studies, (ii) the fact that many cancer types do not have a single publicly available dataset, and (iii) the potential for biomarkers, given that even here there were some clear signatures, we urge the community to collect, and deposit, more *O*-glycomics data of this type.

## Discussion

Our goal here has been to establish a comprehensive, user-friendly, and potent platform for the comparative analysis of glycomics data at various levels of resolution. Accessible via wrapper functions from glycowork, we aim to provide experimentalists as well as bioinformaticians with easy access to state-of-the-art tools for analyzing glycomics data. Throughout this work, we have demonstrated that the workflows we developed for this are fast, robust, sensitive, and rigorous. We are also enthusiastic to note that our presented workflow yielded numerous results from re-analyzing glycomics datasets for which the original authors either did not detect significant effects or for which most to all reported effects became non-significant after correcting for multiple testing. This showcases the gains that can be made in analyzing glycomics data with a dedicated workflow, despite the relatively modest sample sizes which are commonly collected today.

We note that, while we designed this workflow for analyzing glycomics data with structural resolution (linkages etc.) obtained from typical MS/MS glycomics data, *get_differential_expression* and all other functions also can be used if glycans have only been assigned at the compositional level (or even just the *m/z* level). Except for the motif functionality, all other benefits presented herein would then still be applicable. The same applies to glycan nomenclatures. Glycowork, and our workflow by extension, was designed for glycans written in the IUPAC-condensed nomenclature. Any other nomenclature can still be analyzed on the sequence level, and both glycowork-internal (*canonicalize_iupac*) as well as external (glypy^51^) functionalities allow for nomenclature conversion to analyze glycans at the motif level as well.

While we consider sequence motifs in this work, in principle our workflows are also amenable for analyzing glycans that are featurized in another manner. Examples of this include extracting atomic features of glycans (e.g., charge distribution), which can be retrieved via glyLES^52^, or graph features of glycans (e.g., betweenness centrality), which can be calculated within glycowork. As glyLES is also supported within glycowork, all these, and other conceivable featurization methods, provide future opportunities to further explore differential expression of glycan features.

While the work presented here focuses on the analysis of glycomics data, we envision that appropriate data transformations could also make data from glycoproteomics amenable to analysis by the herein developed workflow, yielding insights into system-wide changes in glycosylation.

One part of the difficulty in analyzing glycomics data lies in the scarcity of data, usually caused by the time-intensive analysis of mass spectrometry data, particularly from heterogeneous cancer data. We envision that efforts to automate this analysis bottleneck, such as our recently presented CandyCrunch platform^15^, will lead to an influx of glycomics data, which will not only further enhance the usefulness of the herein presented methods but also facilitate more meta-analyses and conclusions of common dysregulations in the future.

As a corollary to this point, we are also optimistic that future improvements of the methods presented in this work could include support for more elaborate experimental designs. Our current main differential expression analysis only allows for two defined groups of comparisons, whereas in principle users could be interested in more sophisticated features such as including interaction terms or mixed-effects models. While this is established in fields such as transcriptomics^53, 54^, we concluded that the currently existing data acquisition practices in glycomics place this beyond the scope of the current work but remain optimistic that future methodological improvements will facilitate more complex studies.

## Acknowledgments

This work was funded by a Branco Weiss Fellowship – Society in Science awarded to D.B., by the Knut and Alice Wallenberg Foundation, and the University of Gothenburg, Sweden.

## Author Contributions

D.B., J.L., and J.U. conceived the method. D.B., J.L., and J.U. performed computational analyses. D.B., J.L., and J.U. curated the datasets for this study. D.B., J.L., and J.U. prepared the figures. D.B. supervised. D.B. acquired the funding for this study. All authors wrote and edited the manuscript.

## STAR Methods

### Resource availability

#### Lead contact

Further information and requests for resources should be directed to and will be fulfilled by the

Lead Contact, Daniel Bojar.

### Materials availability

This study did not generate new unique reagents.

### Data and Code Availability

- Data curated or generated here can be found in the supplementary tables as well as stored as internal datasets within glycowork.
- All relevant code is integrated into glycowork (version 0.8).
- Any additional information required to reanalyze the data reported in this paper is available from the lead contact upon request.

### Method details

#### Data simulation

Working under the assumption of normalized glycomics data, in which all glycans of a sample sum up to a relative abundance of 100 and only positive abundances are permitted, we modelled this type of data via a Dirichlet distribution, as a specific case of a multivariate distribution in which the components (features) are positive and sum to a constant (100 in this case). Our default simulation set-up included 118 glycans, for which we used concentration parameters according to the experimental relative abundances of the first ‘healthy’ replicate from the *N*-glycan dataset found in Ashwood et al.^55^ (Table S4 in Ashwood et al.; Supplementary Table 2 here). Ground truth effects were added by duplicating a subset (default n = 10 per group of effects, i.e., up- or downregulation) of abundances at random, while scaling the concentration parameter of the test condition by the desired effect magnitude. In each run, random glycan structures were sampled from SugarBase^11^ within glycowork and assigned to the simulated abundances. Random sialylated structures were assigned to the group of upregulated glycans, while random fucosylated structures were assigned to the group of downregulated glycans.

For evaluating the motif-level workflow, the abundance of each glycan containing the motif of interest was scaled by 1.25.

For evaluating imputation strategies, we varied the proportion of randomly missing values from 0 to 70% in ten linearly spaced steps, using a concentration parameter scaling factor of 5.

For evaluating normalization strategies, we varied the concentration parameter scaling, a measure of simulated effect strength, from 1 to 10 in seven linearly spaced steps.

The stability of the simulation results was evaluated by performing the imputation strategy test using relative abundances from different experimental datasets as the concentration parameters for the Dirichlet distribution (Fig. S5)

#### Dataset collection

All glycomics datasets were gathered from the academic literature and were present as supplementary tables in the original articles. We uniformly formatted all these tables for our analyses and they can be found in Supplementary Table 1. Glycan sequences were reformatted into IUPAC-condensed via the *canonicalize_iupac* function from glycowork^11^, followed by a manual inspection. Only glycans where at least a defined core topology (including floating substituents) was annotated were considered for our analyses. We reiterate that this is not necessary for sequence-level analysis with our presented methods but, as we also wanted to explore motif- and motif set-level analyses, we focused on uniform data standards.

For the meta-analysis, we used PubMed, Google Scholar, and Google to search for relevant articles with keyword combinations such as “cancer+glycome” or “tumor+glycome”. All articles were screened and only articles with (i) *O*-glycomics data, (ii) data from tumors, (iii) data from healthy controls, (iv) at least two samples per condition, and (v) at least defined glycan topologies were retained. Then, these articles were processed as described above. The resulting data can be found in Supplementary Table 1.

#### Data normalization

As a first step, we removed variables which had missing values in more than half the replicates, with the rest of the missing values being subject to data imputation described below. We note that this initial removal step can be customized with a keyword argument (‘min_samples’, specifying how many replicates per group need to exhibit a nonzero value for retaining the glycan). The standard normalization procedure employed here was division of each abundance by the total abundance of a sample, essentially normalizing to a total of 100 per sample. Variance stabilizing normalization, used prior to statistical comparisons, included log-transformation of the data and standard scaling to zero mean and unit variance.

#### Data imputation

Data imputation proceeded via the MissForest^30^ approach: all missing values were recorded and then first replaced by the sample median. Then, for each sample, a Random Forest Regressor (default parameters as defined in scikit-learn, version 1.2.2) was trained on all other samples to predict the values at the positions holding missing values. This imputed dataframe was then used in another round of model training and prediction, for a total of five iterations. Finally, we added a small constant (1e-6) to ensure that no zero values were present in the final dataset.

#### Motif extraction and quantification

Motifs were extracted and quantified at three different levels, informed by domain knowledge and accessible via keywords to the *get_differential_expression* function within glycowork, which can be freely combined.

*known*: A set of 154 named motifs (e.g., “LewisX”), defined within glycowork, is used to count the number of occurrences within each glycan via subgraph isomorphism tests. Defined monosaccharide positions (terminal/internal/flexible) ensure matching of structures at appropriate positions (e.g., “Oglycan_core1” (i) matches only at the reducing end and (ii) does not double-count core 2 structures as core 1).

*terminal*: For all non-reducing termini, the monosaccharide and its linkage are extracted and all unique combinations are counted in each glycan. Then, in an effort of feature engineering, linkage uncertainties are tentatively introduced (e.g., “Neu5Ac(a2-?)” if “Neu5Ac(a2-3)” is present) to probe whether they result in new/additional information compared to their defined versions. If not (i.e., if they result in identical results), they are removed again. The purpose here is to capture broader enrichments (e.g., enriched in the umbrella term “Neu5Ac(a2-?)”, versus weaker enrichments in the more specified “Neu5Ac(a2-3)” or “Neu5Ac(a2-6)”).

*exhaustive*: All occurring mono- and disaccharide motifs in the provided glycans are counted for each glycan. Similar to the procedure described above for ‘terminal’, we tentatively introduced uncertainties (e.g., “Gal(b1-?)GlcNAc”) to capture broader effects and trends, yet only retained them if they provided additional information.

We then used these counts to arrive at motif relative abundances by forming a weighted sum for each motif. This was achieved by multiplying the relative abundance of each glycan by its motif count, effectively arriving at the representation of how dominantly exposed a given motif is in a sample.

Next, we automatically deduplicated this quantification via the *clean_up_heatmap* function, as sometimes nominally different motifs (e.g., “Neu5Ac”, “Neu5Ac(a2-3)”, “Neu5Ac(a2-3)Gal”) can have the same relative abundance distribution (e.g., if the only occurrence of “Neu5Ac” in a dataset is within “Neu5Ac(a2-3)Gal”). This was done to reduce the number of statistical tests and hence increase the sensitivity of our analyses. If multiple motifs resulted in the exact same distribution, only one was retained, with the prioritization of named motif > disaccharide > terminal > monosaccharide, guided by the principle of identifying the largest substructure that exhibits this enrichment, which should ease interpretability.

#### Deriving motif abundances from GlyCompare

GlyCompare was implemented via GlyCompareCT^56^. As the recommended conda installation could not be created in our hands, an alternative approach was taken by manually installing certain packages from the environment list in a new conda environment. First, the GlyCompare version was specified as git+https://github.com/LewisLabUCSD/GlyCompare.git@15b415b. Then, other packages were installed at their required specific versions: pandas==1.3.5, bootstrapped==0.0.2, lxml==4.7.1, scipy==1.7.3, and networkx==2.4. The basal cell carcinoma data^26^ then was converted into GlycoCT using the structure translation API from GlyConnect^57^. Once the structural annotation file was prepared along with the original abundance file, as per the GlycompareCT ‘Prepare data’ Step 1a and 1b, GlycompareCT.py was run using the default parameters. The motif_annotation.csv file, mapping the motif codes to WURCS, was then used to convert the motifs into IUPAC-condensed with the GlyCosmos GlycanFormatConverter^58^. The motif abundance table with converted nomenclature was then analyzed using *get_differential_expression* with motifs=False.

#### Enrichment analysis

To form automated sets that were tested for differential expression via multivariate comparisons, we calculated a pairwise correlation matrix from either sequences or motifs. Then, an adjacency matrix was built by forming a link between two sequences/motifs, if and only if they positively correlated above a set threshold (Pearson’s correlation coefficient; default in this work: 0.9) but below 1. Connected components within the graph described by this adjacency matrix were then used as glycan/motif sets for multivariate comparisons.

#### Differential expression analysis

Using variance-based filtering, all sequences/motifs below a minimum variance cut-off value (default in this work: 0.01, or 1%) were removed before differential expression analysis. Then, the data were split into the sample groups defined for comparison. Log-fold changes were calculated as the log_2_-transformed ratio of group means. In the multivariate scenario, log-fold changes of all set-members were averaged. Then, variance stabilization normalization, with the group-specific means and standard deviations, was used before statistical testing. For the univariate case, two-tailed Welch’s t-tests were used to compare normalized abundances, while Hotelling’s T^2^ test was used in the multivariate case^28^. Paired samples are also supported within *get_differential_expression*, using paired t-tests and Cohen’s *d_z_* in the univariate setting^59^. For each comparison, we then further calculated an effect size, Cohen’s *d* for univariate comparisons and the Mahalanobis distance for multivariate comparisons^29^. Further, in all settings, we also used Levene’s test for equality of variances. In any scenario, all resulting p-values were corrected for multiple testing by the Benjamini-Hochberg procedure.

#### ANOVA

Data processing for ANOVA analyses, via the *get_glycanova* function, mirrored those of differential expression analysis above. For each feature, glycan or motif, a linear model was fitted on the abundance via ordinary least squares regression. This was followed by generating an ANOVA table and the corresponding F statistics. Resulting p-values were corrected for multiple testing by the Benjamini-Hochberg procedure. If any ANOVA result reached the significance threshold of 0.05, a Tukey’s HSD (honestly significant difference) test was triggered for all pairwise comparisons of this glycan or motif. Adjusted p-values of lower than 0.05 were then retained as significant pairwise differences.

#### Time-series analysis

Data with multiple timepoints were analyzed with the *get_time_series* function, which again exhibited the same data imputation and motif quantification capabilities as the *get_differential_expression* function. Then, for each glycan or motif, an ordinary least squares model was fitted, aiming to explain glycan abundance as a function of time, with a constant intercept. This was followed by t-tests ascertaining whether the slope of the resulting fit was significantly different from zero. If degree > 1 was chosen, a polynomial function of the chosen degree was instead fitted to the data. Then, a one-way F-test was performed to test whether the fitted model significantly reduced the residuals compared to an intercept-only model. Resulting p-values were corrected for multiple testing by the Benjamini-Hochberg procedure.

#### Meta-analysis via fixed-effects and random-effects models

To calculate effect sizes across different studies, we used the *get_meta_analysis* function within glycowork to estimate combined effect sizes via a fixed-effects model. Here, the individual effect sizes were averaged, weighted by the inverse of their variance. We then also calculated a two-tailed p-value for this combined effect size. Within glycowork, effect size variances were calculated within *cohen_d* for Cohen’s *d* and estimated via bootstrapping (N = 1000) for the Mahalanobis distance.

For random-effects models, between-study variance was estimated as τ² by the DerSimonian and Laird method^60^ in *get_meta_analysis*. If a filepath was specified, effect sizes for a given sequence of motifs were plotted in Forest plots across studies with their 95% confidence interval.

For the meta-analysis presented in this work, we used the *get_differential_expression* function to analyze all cancer datasets individually on the motif level (feature_set = [‘known’, ‘exhaustive’, ‘terminal’]). If the data came from paired samples (e.g., tumor tissue and adjacent healthy tissue from the same individual), we set paired=True when analyzing the data. We then collected the effect sizes and effect size variances for each motif and, for each motif, called the *get_meta_analysis* function with model=’fixed’ to calculate a combined effect size and resulting p-value. Then, all p-values were corrected for multiple testing via the Benjamini-Hochberg procedure.

### Quantification and statistical analysis

For statistical analysis, this study used two-tailed Welch’s t-test for univariate and Hotelling’s T^2^ test for multivariate comparisons. Differences in variance were tested by Levene’s test. Pairwise post-hoc comparisons were done with Tukey’s HSD (honestly significant difference) test. All multiple testing corrections were done via the Benjamini-Hochberg procedure. Effect sizes were estimated via Cohen’s *d* / *d_z_* for univariate and the Mahalanobis distance for multivariate comparisons. All statistical testing has been done in Python 3.11.3 using the glycowork package (version 0.8), the statsmodels package (version 0.14) and the scipy package (version 1.11). Data normalization and motif quantification was done with glycowork (version 0.8).

## Supplementary Tables

**Supplementary Table 1. All curated glycomics datasets used for differential expression analysis.** Related to STAR Methods.

**Supplementary Table 2. Experimental glycan abundances used to establish glycomics simulations.** Related to Figure STAR Methods.

**Supplementary Table 3. Differential expression results using motif abundances gained from GlyCompare or glycowork.** Related to Figure 3.

**Supplementary Table 4. Normalized motif abundances and results from the time series analysis of *N*- and *O*-glycomics data of macrophage differentiation.** Related to Figure 4.

**Supplementary Table 5. Combined effect sizes and p-values for *O*-glycan sequences and motifs from the cancer glycomics meta-analysis.** Related to Figure 5.

## References

1. Ruhaak, L. R., Xu, G., Li, Q., Goonatilleke, E. & Lebrilla, C. B. Mass Spectrometry Approaches to Glycomic and Glycoproteomic Analyses. Chem. Rev. 118, 7886–7930 (2018).

2. Varki, A. Biological roles of glycans. Glycobiology 27, 3–49 (2017).

3. National Research Council (US) Committee on Assessing the Importance and Impact of Glycomics and Glycosciences. Transforming Glycoscience: A Roadmap for the Future. (National Academies Press, 2012). doi:10.17226/13446.

4. Julien, S. et al. Selectin Ligand Sialyl-Lewis x Antigen Drives Metastasis of Hormone-Dependent Breast Cancers. Cancer Res. 71, 7683–7693 (2011).

5. Dall’Olio, F., Pucci, M. & Malagolini, N. The Cancer-Associated Antigens Sialyl Lewisa/x and Sda: Two Opposite Faces of Terminal Glycosylation. Cancers 13, 5273 (2021).

6. Hu, M., Lan, Y., Lu, A., Ma, X. & Zhang, L. Glycan-based biomarkers for diagnosis of cancers and other diseases: Past, present, and future. in Progress in Molecular Biology and Translational Science vol. 162 1–24 (Elsevier, 2019).

7. Hayes, C. A., Nemes, S. & Karlsson, N. G. Statistical analysis of glycosylation profiles to compare tissue type and inflammatory disease state. Bioinformatics 28, 1669–1676 (2012).

8. Zhou, Y. & Neelamegham, S. Comparative Glycomics Analysis of Mass Spectrometry Data. In Glycosylation (ed. Davey, G. P.) vol. 2370 97–113 (Springer US, 2022).

9. Bao, B. et al. Correcting for sparsity and interdependence in glycomics by accounting for glycan biosynthesis. Nat. Commun. 12, 4988 (2021).

10. Bojar, D. et al. A Useful Guide to Lectin Binding: Machine-Learning Directed Annotation of 57 Unique Lectin Specificities. ACS Chem. Biol. acschembio.1c00689 (2022) doi:10.1021/acschembio.1c00689.

11. Thomès, L., Burkholz, R. & Bojar, D. Glycowork: A Python package for glycan data science and machine learning. Glycobiology cwab067 (2021) doi:10.1093/glycob/cwab067.

12. Lundstrøm, J., Urban, J., Thomès, L. & Bojar, D. GlycoDraw: a python implementation for generating high-quality glycan figures. Glycobiology cwad063 (2023) doi:10.1093/glycob/cwad063.

13. Coff, L., Chan, J., Ramsland, P. A. & Guy, A. J. Identifying glycan motifs using a novel subtree mining approach. BMC Bioinformatics 21, 42 (2020).

14. de Haan, N. et al. Developments and perspectives in high-throughput protein glycomics: enabling the analysis of thousands of samples. Glycobiology 32, 651–663 (2022).

15. Urban, J., et al. Predicting glycan structure from tandem mass spectrometry via deep learning. bioRxiv (2023).

16. Thomès, L. & Bojar, D. The Role of Fucose-Containing Glycan Motifs Across Taxonomic Kingdoms. Front. Mol. Biosci. 8, 755577 (2021).

17. Thomès, L., Karlsson, V., Lundstrøm, J. & Bojar, D. Mammalian milk glycomes: Connecting the dots between evolutionary conservation and biosynthetic pathways. Cell Rep. 42, 112710 (2023).

18. Madunić, K. et al. Specific (sialyl-)Lewis core 2 *O*-glycans differentiate colorectal cancer from healthy colon epithelium. Theranostics 12, 4498–4512 (2022).

19. Robbe-Masselot, C. et al. Expression of a Core 3 Disialyl-Le ^x^ Hexasaccharide in Human Colorectal Cancers: A Potential Marker of Malignant Transformation in Colon. J. Proteome Res. 8, 702–711 (2009).

20. Mereiter, S. et al. Glycomic and sialoproteomic data of gastric carcinoma cells overexpressing ST3GAL4. Data Brief 7, 814–833 (2016).

21. Adamczyk, B. et al. Sample handling of gastric tissue and O-glycan alterations in paired gastric cancer and non-tumorigenic tissues. Sci. Rep. 8, 242 (2018).

22. Fernandes, E. et al. Nucleolin-Sle A Glycoforms as E-Selectin Ligands and Potentially Targetable Biomarkers at the Cell Surface of Gastric Cancer Cells. Cancers 12, 861 (2020).

23. Jin, C. et al. Structural Diversity of Human Gastric Mucin Glycans. Mol. Cell. Proteomics 16, 743–758 (2017).

24. Hinneburg, H. et al. Unlocking Cancer Glycomes from Histopathological Formalin-fixed and Paraffin-embedded (FFPE) Tissue Microdissections. Mol. Cell. Proteomics 16, 524–536 (2017).

25. Kawahara, R. et al. The Complexity and Dynamics of the Tissue Glycoproteome Associated With Prostate Cancer Progression. Mol. Cell. Proteomics 20, 100026 (2021).

26. Möginger, U. et al. Alterations of the Human Skin N- and O-Glycome in Basal Cell Carcinoma and Squamous Cell Carcinoma. Front. Oncol. 8, 70 (2018).

27. Amara, A. et al. Networks and Graphs Discovery in Metabolomics Data Analysis and Interpretation. Front. Mol. Biosci. 9, 841373 (2022).

28. Hotelling, H. The Generalization of Student’s Ratio. Ann. Math. Stat. 2, 360–378 (1931).

29. Del Giudice, M. Heterogeneity Coefficients for Mahalanobis’ *D* as a Multivariate Effect Size. Multivar. Behav. Res. 52, 216–221 (2017).

30. Stekhoven, D. J. & Bühlmann, P. MissForest—non-parametric missing value imputation for mixed-type data. Bioinformatics 28, 112–118 (2012).

31. Harris, L., Fondrie, W. E., Oh, S. & Noble, W. S. Evaluating proteomics imputation methods with improved criteria. http://biorxiv.org/lookup/doi/10.1101/2023.04.07.535980 (2023) doi:10.1101/2023.04.07.535980.

32. Rasch, D., Kubinger, K. D. & Moder, K. The two-sample t test: pre-testing its assumptions does not pay off. Stat. Pap. 52, 219–231 (2011).

33. Lumley, T., Diehr, P., Emerson, S. & Chen, L. The Importance of the Normality Assumption in Large Public Health Data Sets. Annu. Rev. Public Health 23, 151–169 (2002).

34. Tsagris, M., Alenazi, A., Verrou, K.-M. & Pandis, N. Hypothesis testing for two population means: parametric or non-parametric test? J. Stat. Comput. Simul. 90, 252–270 (2020).

35. Terra Machado, D., Bernardes Brustolini, O. J., Côrtes Martins, Y., Grivet Mattoso Maia, M. A. & Ribeiro De Vasconcelos, A. T. Inference of differentially expressed genes using generalized linear mixed models in a pairwise fashion. PeerJ 11, e15145 (2023).

36. Dagogo-Jack, I. & Shaw, A. T. Tumour heterogeneity and resistance to cancer therapies. Nat. Rev. Clin. Oncol. 15, 81–94 (2018).

37. Sibille, E. et al. Ganglioside Profiling of the Human Retina: Comparison with Other Ocular Structures, Brain and Plasma Reveals Tissue Specificities. PLOS ONE 11, e0168794 (2016).

38. Hinneburg, H. et al. High-resolution longitudinal N- and O-glycoprofiling of human monocyte-to-macrophage transition. Glycobiology 30, 679–694 (2020).

39. Mohammad, M. A., Hadsell, D. L. & Haymond, M. W. Gene regulation of UDP-galactose synthesis and transport: potential rate-limiting processes in initiation of milk production in humans. Am. J. Physiol.-Endocrinol. Metab. 303, E365–E376 (2012).

40. Dettori, J. R., Norvell, D. C. & Chapman, J. R. Fixed-Effect vs Random-Effects Models for Meta-Analysis: 3 Points to Consider. Glob. Spine J. 12, 1624–1626 (2022).

41. Chatterjee, S. et al. Protein Paucimannosylation Is an Enriched *N*-Glycosylation Signature of Human Cancers. PROTEOMICS 19, 1900010 (2019).

42. Munkley, J. & Elliott, D. J. Hallmarks of glycosylation in cancer. Oncotarget 7, 35478–35489 (2016).

43. Brockhausen, I. Mucin-type *O*-glycans in human colon and breast cancer: glycodynamics and functions. EMBO Rep. 7, 599–604 (2006).

44. Chugh, S. et al. Pathobiological implications of mucin glycans in cancer: Sweet poison and novel targets. Biochim. Biophys. Acta BBA - Rev. Cancer 1856, 211–225 (2015).

45. Steentoft, C. et al. Glycan-directed CAR-T cells. Glycobiology 28, 656–669 (2018).

46. Walker, M. R. et al. O-linked α2,3 sialylation defines stem cell populations in breast cancer. Sci. Adv. 8, eabj9513 (2022).

47. Tanaka-Okamoto, M. et al. Various sulfated carbohydrate tumor marker candidates identified by focused glycomic analyses. Glycobiology glycob;cww133v2 (2016) doi:10.1093/glycob/cww133.

48. Dobie, C. & Skropeta, D. Insights into the role of sialylation in cancer progression and metastasis. Br. J. Cancer 124, 76–90 (2021).

49. Munkley, J. The Role of Sialyl-Tn in Cancer. Int. J. Mol. Sci. 17, 275 (2016).

50. Ju, T., Otto, V. I. & Cummings, R. D. The Tn Antigen-Structural Simplicity and Biological Complexity. Angew. Chem. Int. Ed. 50, 1770–1791 (2011).

51. Klein, J. & Zaia, J. glypy: An Open Source Glycoinformatics Library. J. Proteome Res. 18, 3532–3537 (2019).

52. Joeres, R., Bojar, D. & Kalinina, O. V. GlyLES: Grammar-based Parsing of Glycans from IUPAC-condensed to SMILES. J. Cheminformatics 15, 37 (2023).

53. Hoffman, G. E. & Roussos, P. Dream: powerful differential expression analysis for repeated measures designs. Bioinformatics 37, 192–201 (2021).

54. Rivellese, F. et al. Rituximab versus tocilizumab in rheumatoid arthritis: synovial biopsy-based biomarker analysis of the phase 4 R4RA randomized trial. Nat. Med. 28, 1256–1268 (2022).

55. Ashwood, H. E. et al. Characterization and statistical modeling of glycosylation changes in sickle cell disease. Blood Adv. 5, 1463–1473 (2021).

56. Zhang, Y. et al. Preparing glycomics data for robust statistical analysis with GlyCompareCT. STAR Protoc. 4, 102162 (2023).

57. Alocci, D. et al. GlyConnect: Glycoproteomics Goes Visual, Interactive, and Analytical. J. Proteome Res. 18, 664–677 (2019).

58. Tsuchiya, S., Yamada, I. & Aoki-Kinoshita, K. F. GlycanFormatConverter: a conversion tool for translating the complexities of glycans. Bioinformatics 35, 2434–2440 (2019).

59. Lakens, D. Calculating and reporting effect sizes to facilitate cumulative science: a practical primer for t-tests and ANOVAs. Front. Psychol. 4, (2013).

60. DerSimonian, R. & Laird, N. Meta-analysis in clinical trials. Control. Clin. Trials 7, 177–188 (1986).

