## Supplemental Figures for "Decoding Glycomics: Differential Expression Reimagined"

### Supplementary Figures

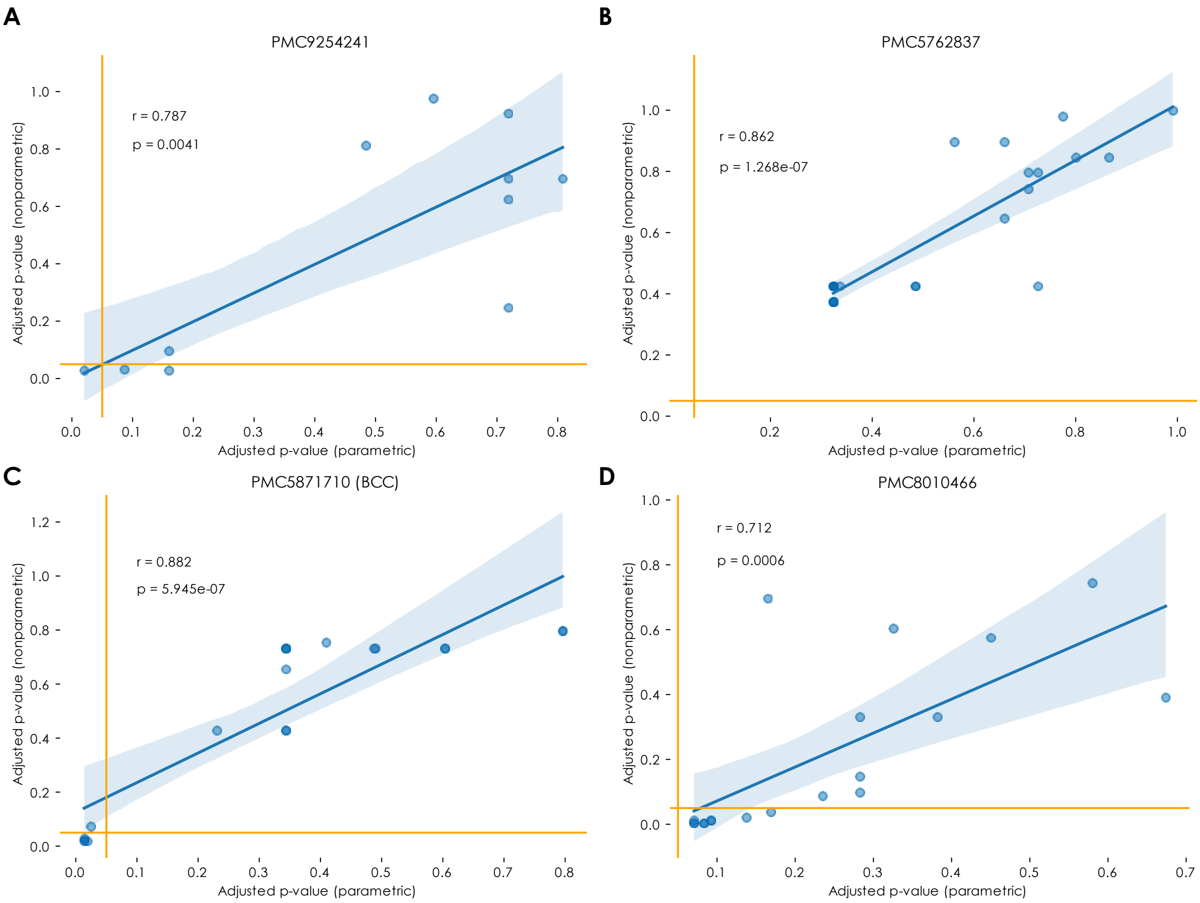


**Figure S1. Comparing parametric and nonparametric tests for differential glycomics expression analysis; Related to Figure 2A. A-D)** For four example glycomics datasets, PMC9254241 (A), PMC5762837 (B), the BCC dataset within PMC5871710 (C), and PMC8010466 (D), we used our *get_differential_expression* pipeline for all motifs, with the only difference being using a parametric Welch's t-test versus a nonparametric Mann-Whitney U test for statistical testing. Shown are the adjusted p-values of both approaches for each motif, fitted with an ordinary least squares regression (shown with its 95% confidence band). Model fit is reported by Pearson's r as a correlation coefficient, and a p-value for the t-test of the regression coefficient against zero. Orange vertical and horizontal lines indicate conventional p-value cut-offs of 0.05 to establish statistical significance.

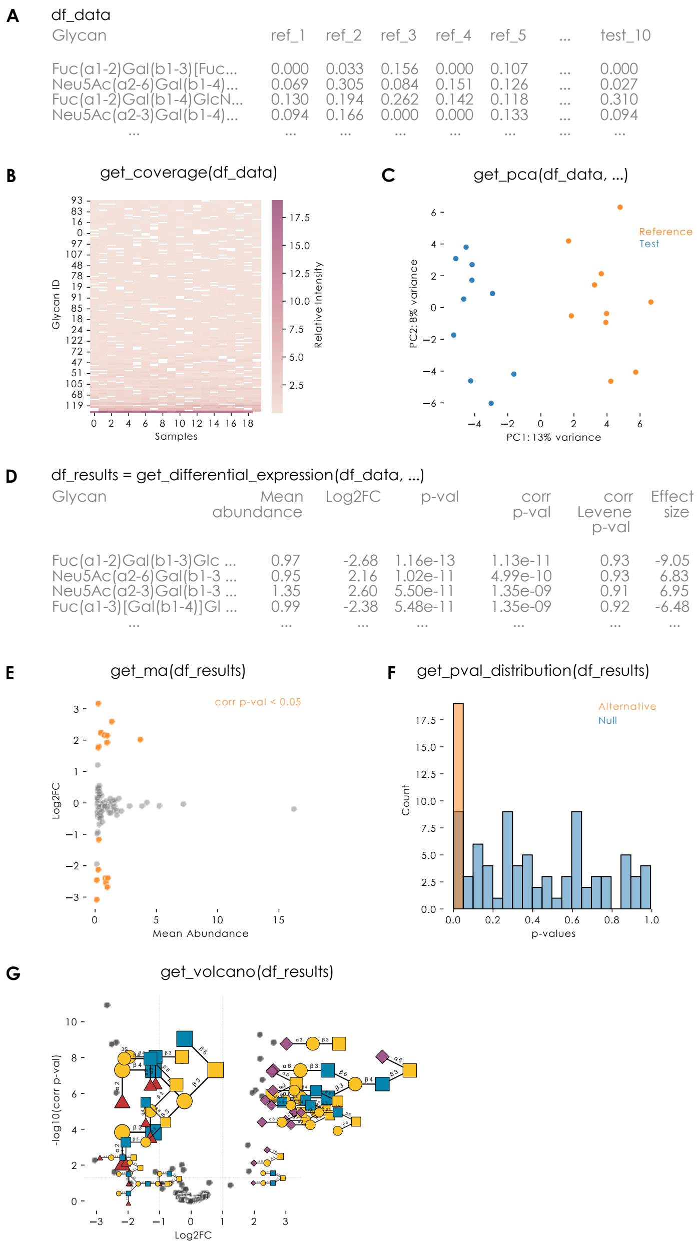


**Figure S2. Full differential expression analysis workflow with example plotting options; Related to Figure 2. A)** Simulated glycomics dataset (see STAR Methods for details), with 10 replicates per condition (reference and test). In the test condition, a group of 10 upregulated, sialylated glycans and a group of 10 downregulated, fucosylated glycans had their concentration parameters multiplied or divided, respectively, by a factor of 5. The dataset included 10% missing values (missing at random). **B)** The *get_coverage* function (*glycowork.motif.analysis*) displays glycan coverage across samples, ordered by average abundance. **C)** The *get_pca* function (*glycowork.motif.analysis*) performs a principal component analysis and plots the two first components, displaying their explained variance in percent. **D)** The *get_differential_expression* (*glycowork.motif.analysis*) function performs the differential expression analysis. Here, the function arguments included *motifs* = *False* (for sequence-level analysis) and *impute* = *True* (for MissForest-based data imputation). **E)** The *get_ma* function (*glycowork.motif.analysis*) returns an MA plot (log2FC vs mean abundance) of the differential expression results. **F)** The *get_pval_distribution* (*glycowork.motif.analysis*) function plots a histogram of the calculated differential expression p-values. **G)** The *get_volcano* (*glycowork.motif.analysis*) plots the differential expression results as -log10(corr p-val) vs log2FC, commonly referred to as a volcano plot. Here, the figure was further annotated using the *annotate_figure* function of GlycoDraw (*glycowork.motif.draw*).

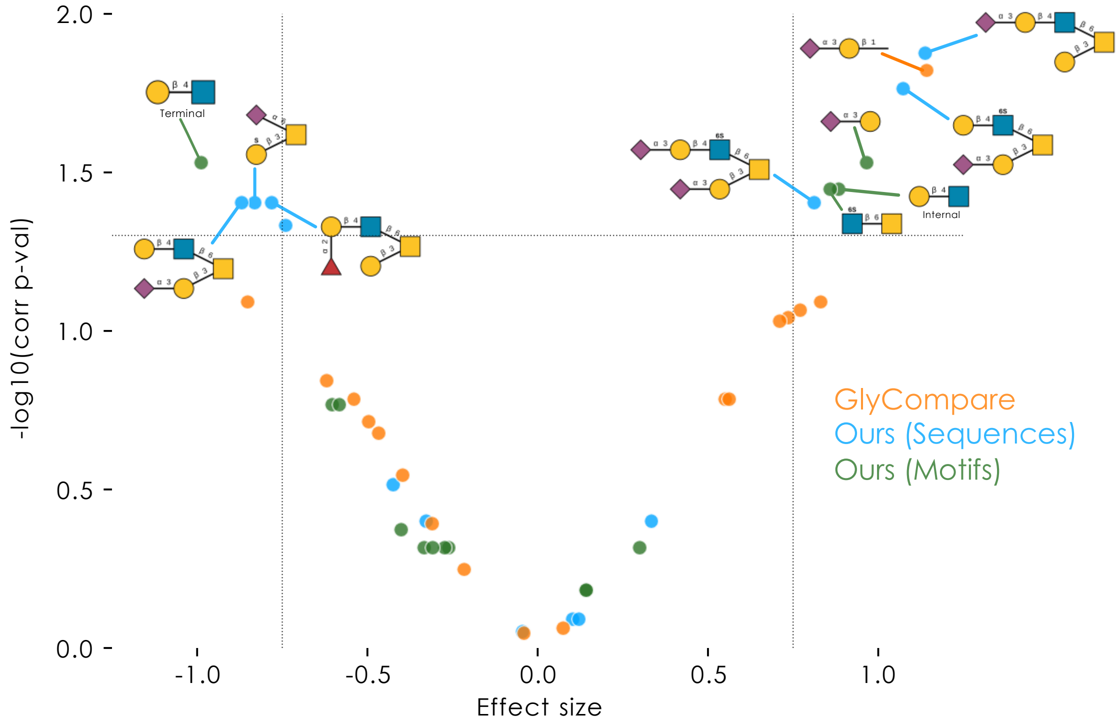


**Figure S3. Comparing motifs derived from GlyCompare and glycowork for differential glycomics expression analysis; Related to Figure 3.** The same FFPE tissue *O*-glycomics data as in Fig. 2F was analyzed using *get_differential_expression* with motifs=False in blue and motifs=True in green, without any further pre-processing. The analysis carried out using motif abundances generated with GlyCompare (see STAR Methods) with motifs=False is depicted in orange. Results are depicted as overlayed volcano plots created using *get_volcano*. Full results can be found in Supplementary Table 3.

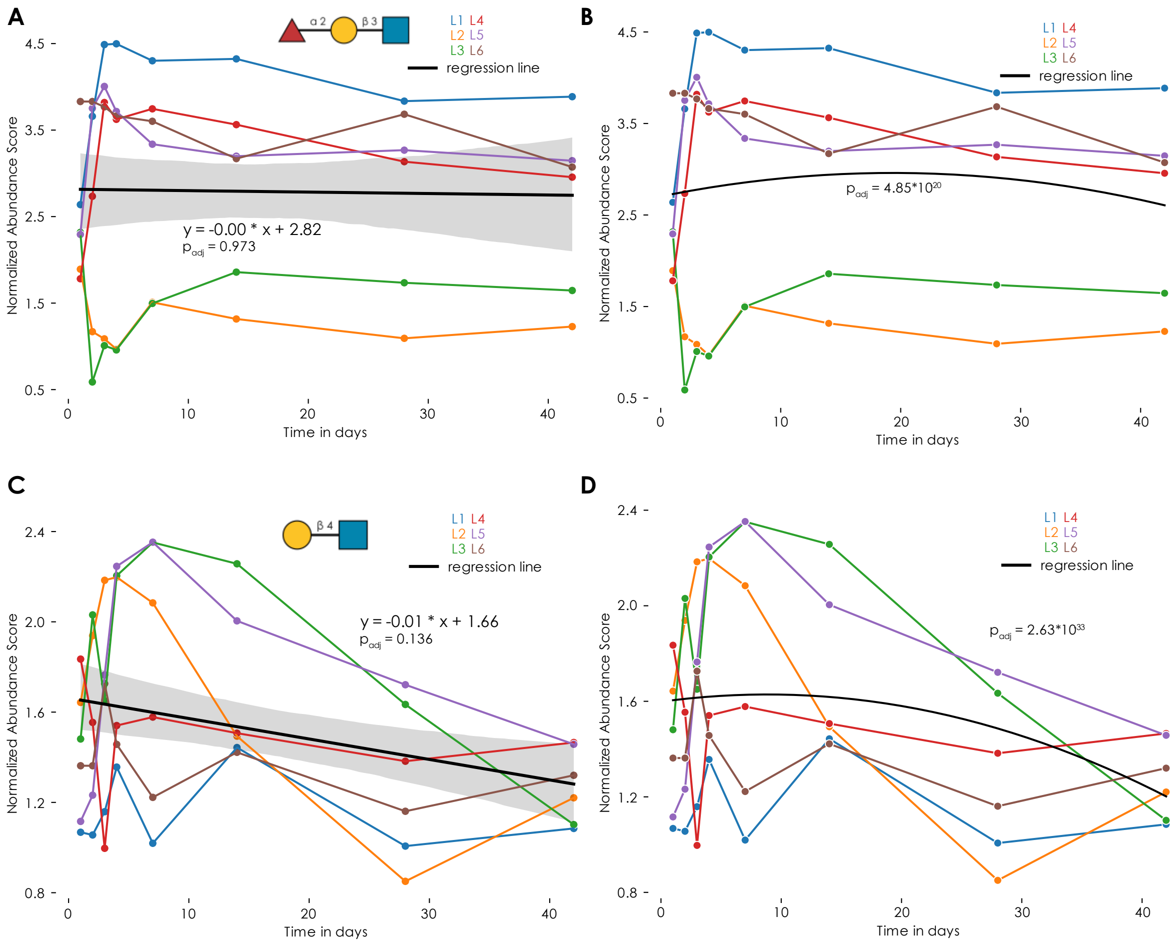


**Figure S4. Fitting nonlinear trends with the *get_time_series* function; Related to Figure 4. A-D)** Using the measured free milk oligosaccharide time series data from Mohammad et al. (doi: 10.1152/ajpendo.00175.2012), we used the *get_time_series* function with degree = 1 (A,C; linear regression) or degree = 2 (B, D; polynomial regression) on the motif level, with the examples of H type 1 (A,B) and LacNAc type 2 (C,D) motifs shown. Curves respond to normalized relative motif abundances of each participant. The p-values are derived from a t-test of the regression coefficient against zero (A,C) or a one-way F-test of whether residuals are significantly reduced compared to an intercept-only model (B,D). All p-values were corrected for multiple testing with the Benjamini-Hochberg procedure.

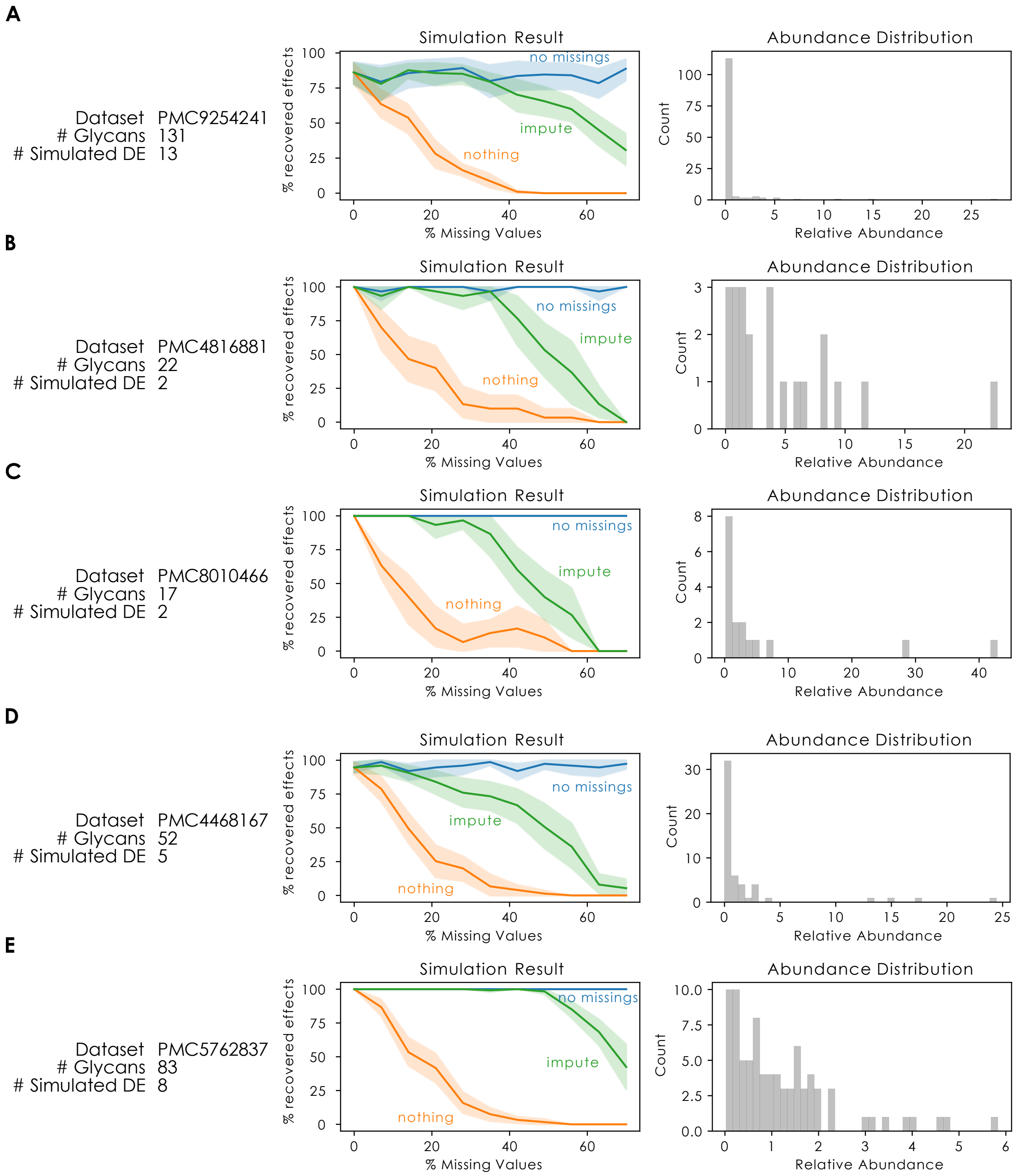


**Figure S5. Evaluating simulation stability to template abundance data; Related to STAR Methods. A-E)** With concentration parameters defined by the relative glycan abundance from various experimental datasets, we simulated glycomics data with 0 to 70% missing values and tested the sensitivity of the imputation workflow as in Figure 2C. For each dataset, ground truth effects were added by duplicating a subset (10% of total, rounded up) of abundances at random, and scaling the concentration parameters by a factor of 5. 15 simulations per level of missing data were performed.
